# Computational Model for Simulating Drug-Induced Arrhythmia Sensitivity of Human iPSC-derived Cardiomyocytes

**DOI:** 10.1101/236240

**Authors:** Xin Gao, Yue Yin, Neil Daily, Tyler Engel, Li Pang, Brian E. Carlson, Tetsuro Wakatsuki

## Abstract

A mathematical model describing the electrophysiology and ion handling of cardiomyocytes is a complement to experimental analysis predicting drug-induced proarrhythmic potential in humans as proposed by Comprehensive *in vitro* Proarrhythmia Assay (CiPA). While CiPA endorses the use of the O’Hara Rudy (ORd) model, which was developed to simulate electrophysiology of human adult ventricular cardiomyocytes (hAVCMs), to predict drug-induced proarrhythmias; the human induced pluripotent stem cell derived-cardiomyocytes (hiPSC-CMs) was proposed for experimental verifications. The hiPSC-CMs, especially cultured in 2D culture dishes, are inherently different from hAVCMs exhibiting different ion channel density and an immature sarcoplasmic reticulum function. To reconcile this mismatch, we have developed a mathematical electrophysiology model of an hiPSC-CM by incorporating differences in gene expressions of ion channels, pumps and receptors in hiPSC-CMs against those found in hAVCMs. This model can be used to model any hiPSC-CM cell line where expression data has been obtained and replaces the background currents for K^+^ and Na^+^ in the ORd model with the known ultra-rapid K^+^ channel and hyperpolarization activated K^+^/Na^+^ channel currents. With this new model, three batches each from two different hiPSC-CM cell lines are compared to experimental data of action potential duration. This mathematical model recapitulates a ventricular-like action potential morphology with a Phase 2 plateau lasting a few hundred milliseconds. However, the resting membrane potential is not as depolarized (−65 to −70 mV) as that of hAVCM (-80 mV). The elevated resting membrane potential matches experimental data previously and is thought to keep rapid sodium channels from triggering membrane depolarization instead being replaced by an overexpression of L-Type Ca^2+^ channel current. This model is a key step in identifying the variability between different hiPSC-CM lines and even batches of the same line, opening the door to realizing analysis of patient specific preparations in the future.

## Introduction

Over 130,000 (~1/3 of total cardiac adverse events) cases of drug induced, potentially life-threatening, cardiac arrhythmias, *torsades de points* (TdP), were recorded by the US Food and Drug Administration (FDA) between 1969 and 2000^1^. Some of these drugs were withdrawn from the market due to torsadogenic risk^2^. After the International Conference on Harmonization (ICH) issued ICHS7B and E14 Guidelines in 2005^3^, the rate of TdP reports has declined^4^. The S7B guideline recommends performing a simple hERG channel inhibition assay using cell lines (e.g., CHO or HEK cells) expressing the hERG channel as the primary *in vitro* assay to evaluate TdP risk. Despite its well-recognized role in assessing drug-induced QTc prolongation risk, solely based on hERG potency and therapeutic plasma exposure can result in incorrected conclusions and unwanted compound attrition during drug discovery^5^.

The Comprehensive *in vitro* Proarrhythmia Assay (CiPA) Initiative has defined a new paradigm by which computational models are identified with experimental data generated with heterologous systems to predict drug-induced proarrhythmia risk in human ventricular myocytes. These proposed theoretical models incorporate current density or flux and dynamic gating function of the major ion channels, pumps and receptors expressed in CMs. In coupling with detailed experimental data generated with hiPSC-CMs, the new *in vitro* assay systems can provide a more comprehensive view of cardiomyocyte function than that relying simply on compound-effects of hERG channel inhibition^6^. hiPSC-CMs express the same ion channels, pumps and receptors known to be expressed in human adult ventricular cardiomyocytes (hAVCMs) however the level of expression has been observed to be different^7^. hiPSC-CMs also exhibit action potentials (AP) profiles similar in form to hAVCMs but with a large variation in action potential duration at 90% repolarization (APD_90_) of 70–789 ms^8^ and spontaneous beating frequencies of 25-85 beats per minute^7^.

While hiPSC-CM derived tissues are a logical choice to improve the guideline^9^, their immaturity and under-developed excitation-contraction (EC) coupling^10,11^ are also of critical concern. A previous theoretical model of hiPSC-CMs electrophysiology and ion handling has been developed^12^ based on experimental data of ion channel function in hiPSC-CMs at 30-53 days post differentiation^13^. While the previous model by Paci et al. is extremely successful in describing purified hiPSC-CMs at the 30-53 day maturation time point, a patch clamp study similar to that performed by Ma et al. would be needed to characterize other maturation time points or variation across different induced pluripotent stem cell lines. Therefore, we took a different approach using the latest hAVCM theoretical model by O’Hara et al.^14^ (ORd) coupled with relative expression data between hAVCMs and hiPSC-CMs for 11 genes associated with ion channel, pump, receptor and transporter density to determine the change of ionic transport driving electrophysiological function in an hiPSC-CM. In this new model, we have replaced the background currents for K^+^ and Na^+^ with the known ultra-rapid K^+^ channel and hyperpolarization activated K^+^/Na^+^ channel currents as shown in Figure 1. This approach for identifying a theoretical model of hiPSC-CM function has three major advantages: 1) relative variations in electrophysiological function can be identified between different batches and cell lines of hiPSC-CMs or different lines of hiPSC-CMs derived from different donors; 2) functions of hiPSC-CMs with different maturation status can be described by utilizing expression data at each maturation time point; and 3) this model can be directly compared to hAVCM function where relative expression is set to one for all ion channels, pumps and transporters. In the methods section, we outline how we have modified the ORd model to simulate electrophysiology of hiPSC-CMs yet preserve the ability for the model upon reparameterization to recapitulate the hAVCM phenotype. In the results below, we will use this model to represent the electrophysiology of three batch variants across two different commercial cell lines where changes in the action potential duration at 90% repolarization (APD_90_) are identified to show qualitative agreement with experimental results. Finally, we predict with this expression-based hiPSC-CM model how concurrent blocking of fast and slow components of the Na^+^ channel, rapid delayed rectifier K^+^ channel and L-type Ca^2+^ channel influences APD_90_ and proarrhythmia potential.

**Figure 1.**
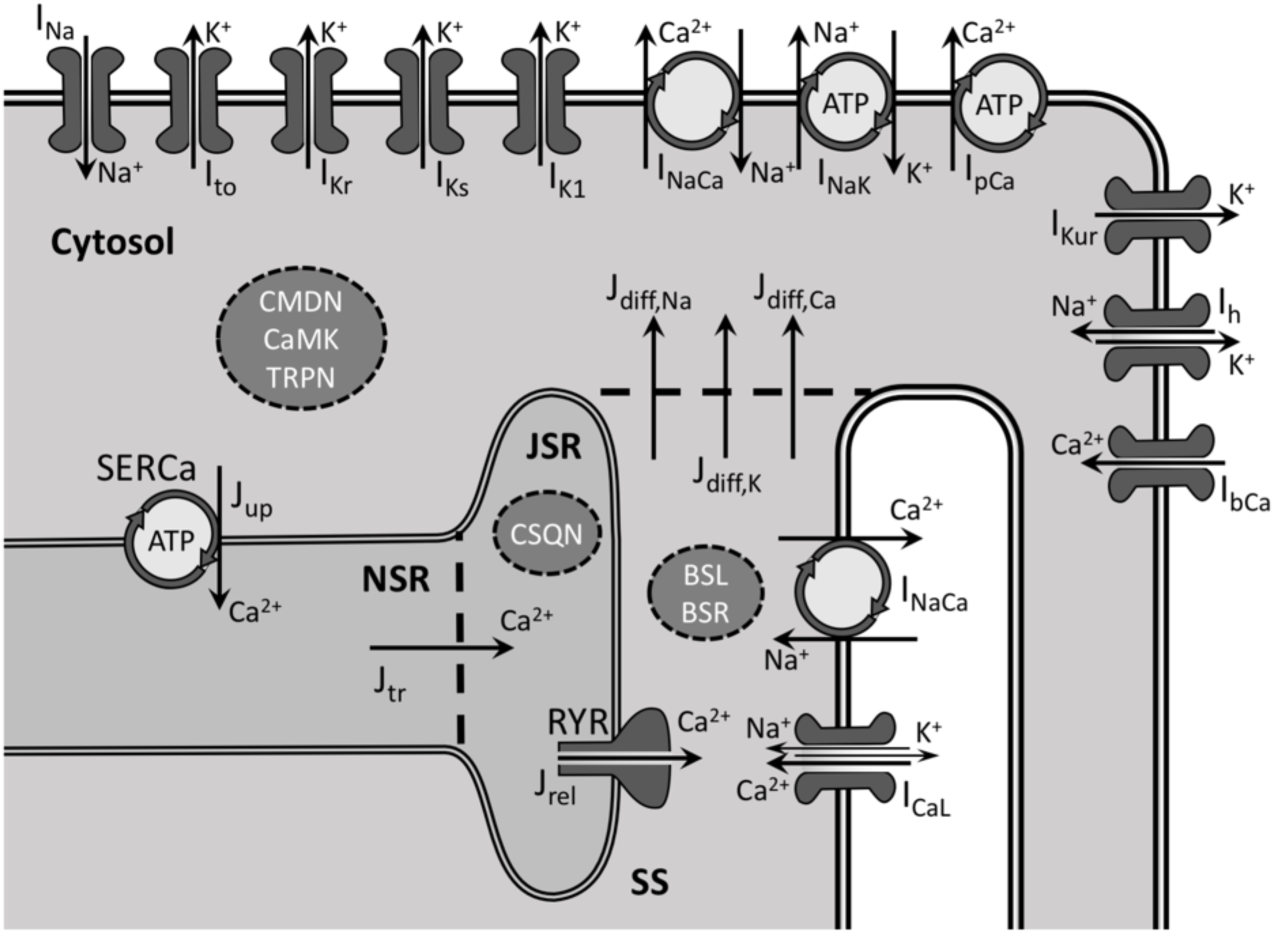
Schematic of the modified ORd model where maximal conductances/permeabilites can be reparameterized to represent the electrophysiology of an hiPSC-CM. Channel, exchanger and pump currents and fluxes are formulated in the same way as in the ORd model: fast Na^+^ (*I_Na_*), transient outward K^+^ (*I_to_*), rapid delayed rectifier K^+^ (*I_Kr_*), slow delayed rectifier K^+^ (*I_Ks_),* inward rectifier K^+^ (I_K1_), Na^+^/Ca^2+^ exchanger (*I_NaCa_*), Na^+^/K^+^ ATPase (*I_NaK_),* plasma membrane Ca^2+^ pump (*I_pCa_),* L-type Ca^2+^ (*I_CaL_),* ryanodine receptor Ca^2+^ (J_rel_), sarcoplasmic/endoplasmic reticulum Ca^2+^ pump (J_up_) and background Ca^2+^ (*I_bCa_).* Two background channel currents are omitted – background Na^+^ and K^+^ – and are replaced with an ultrarapid uptake K^+^ current (*I_Kur_)* and a hyperpolarization-activated Na^+^/K^+^ current (*I_h_*). Buffering of Ca^2+^ is implemented in the same manner as in the ORd model: subspace (SS) buffering by the sarcolemmal and sarcoplasmic reticulum membranes (BSL and BSR, respectively), calmodulin, calmodulin kinase and troponin in the cytosol (CMDN, CaMK and TRPN, respectively) and calsequestrin (CSQN) in the junctional sarcoplasmic reticulum (JSR). Note that the total L-type Ca^2+^ current formulation includes small Na^+^ and K^+^ currents and the total fast Na^+^ current includes both fast and late Na^+^ currents. Na^+^/Ca^2+^ exchangers are distributed across the plasma membrane with 20% in the t-tubules interacting with the SS and 80% in the remaining membrane interacting directly with the cytosol. Diffusional fluxes between the SS and cytosol of all three ions are represented by J_diff,Na_, J_diff_,_Ca_ and J_diff,K_ and Ca^2+^ flux between the network sarcoplasmic reticulum (NSR) and JSR by J_tr_.

## Results

### Simulating a typical action potential morphology of iPSC-CMs

To represent the differences in electrophysiological properties between hiPSC-CMs and hAVCMs, the ORd model was modified to simulate electrophysiology of hiPSC-CMs by scaling conductances and permeabilities with respect to relative gene expressions of the ion channels, exchangers, pumps and receptors (Table 1). It is assumed that the relative gene expression between hAVCMs and hiPSC-CMs of the gene encoding a given channel, exchanger, etc. can be used to linearly scale its associated conductance or permeability in the hiPSC-CM model. Simulations were performed for each of the 3 different batches from two hiPSC-CM lines produced from a commercial supplier of hiPSC-CMs^7^ along with the expression in hAVCM which we assumed to be represented by the ORd model simulated at its published parameters. A simulated AP profile of the hiPSC-CM resembles those obtained experimentally using patch clamp techniques^13^ and membrane potential dyes reflecting APs published previously^15^. Our simulations also indicate that pacing frequency can significantly change morphologies of AP profiles and shows a predicted shorter APD_90_ for Cor4U cell lines than for the iCell lines (Figure 2**a, b**).

**Table 1.** Relative gene expression data for eleven ion channels, pumps, exchangers and receptors in hAVCM and 3 batches from two hiPSC-CM cell lines (iCell and Cor4U). These expression data are from Figure 1 of the study by Huo et al.^7^ where hAVCM is represented by hALV.

| Gene | Ion channel,<br>pump or re-<br>ceptor | Current<br>or Flux | hAVCM | iCell1 | iCell2 | iCell3 | Cor.4U1 | Cor.4U2 | Cor.4U3 |
| --- | --- | --- | --- | --- | --- | --- | --- | --- | --- |
| SCN5A | Fast Na<sup>+</sup> | $I_{Na}$ | 1.000 | 0.656 | 0.765 | 0.777 | 0.371 | 0.391 | 0.303 |
| CACNA1C | L-Type Ca<sup>2+</sup> | $I_{CaL}$ | 1.000 | 4.716 | 4.110 | 5.397 | 3.282 | 2.735 | 1.256 |
| KCNH2 | Rapid Delayed<br>Rectifier K<sup>+</sup> | $I_{Kr}$ | 1.000 | 1.060 | 1.001 | 1.161 | 1.352 | 1.290 | 1.130 |
| KCNQ1 | Slow Delayed<br>Rectifier K<sup>+</sup> | $I_{Ks}$ | 1.000 | 0.987 | 0.976 | 0.965 | 1.100 | 0.966 | 0.892 |
| KCND2 | Transient Out-<br>ward K<sup>+</sup> | $I_{to}$ | 1.000 | 1.464 | 1.902 | 1.660 | 4.893 | 3.096 | 1.267 |
| KCNA5 | Ultra-rapid De-<br>layed Rectifier<br>K<sup>+</sup> | $I_{Kur}$ | 1.000 | 0.0897 | 0.0913 | 0.0620 | 4.415 | 3.851 | 9.876 |
| HCN4 | Hyperpolariza-<br>tion Activated<br>K<sup>+</sup> | $I_f$ | 1.000 | 6.152 | 6.801 | 6.651 | 8.267 | 8.139 | 6.250 |
| KCNJ2 | Inward Recti-<br>fier K<sup>+</sup> | $I_{K1}$ | 1.000 | 0.0555 | 0.0544 | 0.0742 | 0.0946 | 0.0633 | 0.0625 |
| SLC8A1 | Na<sup>+</sup>/Ca<sup>2+</sup> Ex-<br>changer | $I_{NaCa}$ | 1.000 | 1.509 | 1.415 | 1.549 | 2.185 | 2.282 | 1.528 |
| RYR2 | Ryanodine Re-<br>ceptor | $J_{rel}$ | 1.000 | 0.0741 | 0.0768 | 0.108 | 0.210 | 0.135 | 0.262 |
| ATP2A2 | Sarcoplasmic<br>Reticulum Ca<sup>2+</sup><br>Pump | $J_{up}$ | 1.000 | 0.358 | 0.335 | 0.377 | 0.673 | 0.563 | 0.775 |

**Figure 2.**
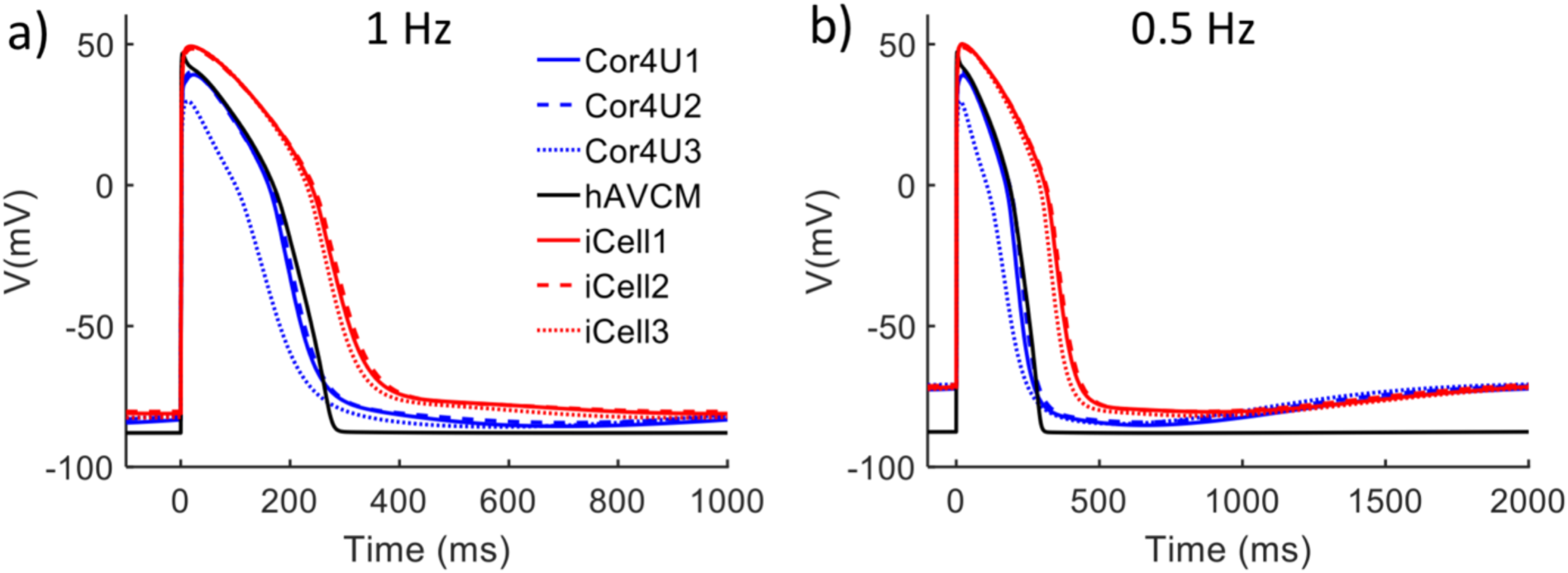
Comparison of simulated action potential profiles generated using gene expression data from two lines of human induced pluripotent stem cell derived cardiomyocytes (hiPSC-CMs). Three different batches of hiPSC-CM from each cell line (Cor4U and iCells) were analyzed and compared with an action potential profile of human adult ventricular cardiomyocytes (hAVCMs) at (**a**) 1 Hz and (**b**) 0.5 Hz.

### Elevated resting membrane potential in hiPSC-CM

As has been previously recorded experimentally^13,14^, hiPSC-CMs have a higher resting membrane potential than that of hAVCMs. Our expression-based hiPSC-CM model, parameterized to represent iCell1, simulated an elevated resting membrane potential of hiPSC-CM to be −81.0 mV at 1 Hz and −71.5 mV at 0.5 Hz, which are both significantly higher than that (−87.9 mV) of hAVCM (Figure 3). Because of the elevated resting membrane potential in hiPSC-CMs the inactivation gate of fast Na^+^ channels do not entirely reset. Therefore, it is posited that the fast Na^+^ channel current, *I_Na_*, does not contribute significantly to the initiation of the hiPSC-CM excitation. Instead an influx of calcium ions through L-type Ca channel, *I_CaL_,* initiates the excitation. Comparison between the simulated *I_Na_* (Figure 3**c, d**) and *I_CaL_* (Figure 3**e, f**) of hAVCM and hiPSC-CM shows an increased *I_CaL_* and reduced *I_Na_* contribution in hiPSC-CMs which were exacerbated further by reducing pacing frequencies to 0.5 Hz. The maximal rate of AP depolarization (dV/dt^max^), of hAVCM and hiPSC-CMs are 342 mV/ms and 134 mV/ms respectively, indicating the hiPSC-CM’s reduced maximum rate of membrane potential depolarization with reduced *I_Na_* contribution. These dV/dt^max^ values are calculated by finding the steepest section of the AP upstroke when the time from AP initiation to peak voltage is divided into 100 sections (see Supplemental Table 1). These simulation dV/dt^max^ values are larger than those reported experimentally^16^ likely because the experimental measures are sampled at a lower frequency. However, the same relative observation can be seen with the dV/dt^max^ of hAVCMs are larger than that of hiPSC-CMs.

**Figure 3.**
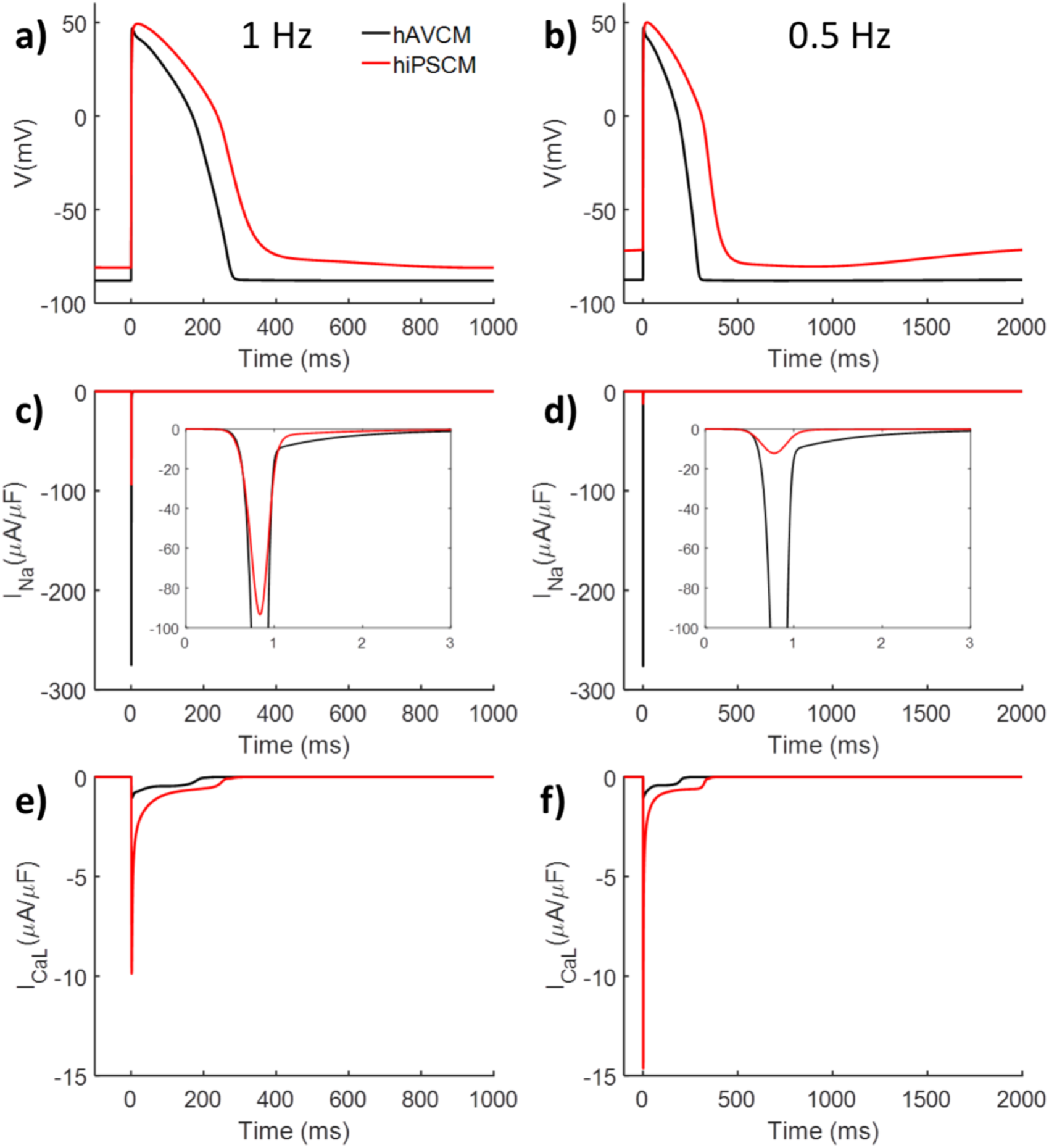
Simulated action potential of hAVCM and hiPSC-CM (iCell1) paced at 1 Hz (**a**) and 0.5 Hz (**b**). Note that diminished fast Na^+^ current at 1 Hz (**c**) and 0.5 Hz (**d**) and enlarged L-type Ca^2+^ current at 1Hz (**e**) and 0.5 Hz (**f**) in hiPSC-CM as compared to hAVCM are shown with the largest discrepancy between hiPSC-CM and hAVCM at 0.5 Hz (**e** and **f**).

### *I_CaL_* and *I_Na_* contributions to electrophysiology of hiPSC-CM and hAVCM

To analyze the extent to which *I_CaL_* and *I_Na_* current influences the shape of AP and calcium transient (CaT) profiles in hiPSC-CMs, we modulated the L-type Ca^2+^ or fast Na^+^ channel conductance in each model with the ORd model representing an hAVCM and the iCell1 as the representative hiPSC-CM (Figure 4). The L-type Ca^2+^ or fast Na^+^ channel conductances were varied in each model from 0 to 5 times that of hAVCM while the remaining ion channels, exchangers, etc. were held constant at the hAVCM and hiPSC-CM values. Without *I_CaL_* current, hiPSC-CM and hAVCM did not exhibit a plateau phase (Figure 4**a, b**). While increasing *I_CaL_* conductance elongated AP duration in hAVCMs and hiPSC-CMs, even a small increment (e.g., *g_CaL_/g_CaL,0_* = 0.5) induced the plateau phase of hiPSC-CM. An increasing Ca^2+^-entry via the *I_CaL_* channel above the level of hAVCM (*g_CaL_/g_CaL,0_* > 1) changed peak of CaT more significantly in hAVCMs than in hiPSC-CMs (Figure 4**c, d**).

**Figure 4.**
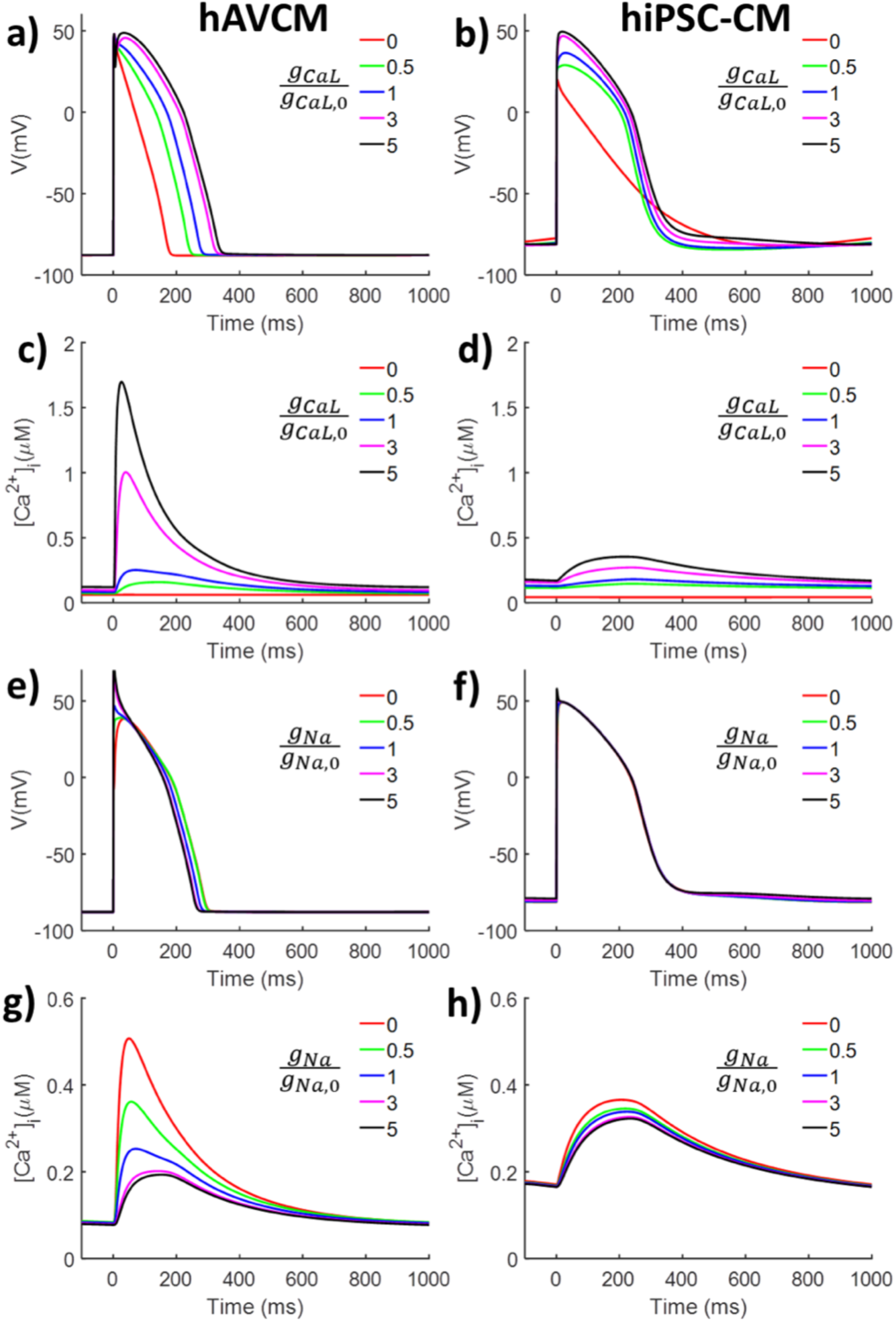
Effects of changing L-type Ca^2+^ channel conductance (*g_CaL_/g_CaL,0_*) on simulated action potential profiles for an hAVCM (**a**) and hiPSC-CM (iCell1) (**b**) and transients of internal calcium concentration ([Ca^2+^]_i_) in (**c**), (**d**) for an hAVCM and hiPSC-CM, respectively. Changing fast Na^+^ channel conductance (*g_Na/_g_Na,0_*) had limited effects on the simulated action potential of hAVCM (**e**) and hiPSC-CM (**f**) as well as on calcium transients of hiPSC-CM (**h**). Changing fast Na^+^ channel conductance (*g_Na/_g_Na,0_*) significantly changed peaks of calcium transients of hAVCM (**g**).

Changing the fast Na^+^ channel conductance had little effects on AP profiles in the hAVCM (Figure 4**e**) but changed amplitudes of CaTs drastically (Figure 4**g**). Even without the *I_Na_* current (*g_Na_/g_Na,0_*= 0), the pacing current (-80 mV pulse for 1 ms) in the simulation was enough to activate *I_CaL_* and generate an AP in the hAVCM. An increasing *I_Na_* current elevated the maximal slope of the membrane potential, (dV/dt)^max^ in the hAVCM from 80 mV/s at *g_Na_/g_Na,0_* = 0 to 435 mV/s at *g_Na_/g_Na,0_* = 5 (sampled at 100 time divisions on upstroke). Conversely, the changes in *I_Na_* current had almost no effect on the hiPSC-CM profiles of AP and a very little effect on CaT (Figure 4**g, h**).

### Modulating *I_K1_* but not *I_h_* restores resting membrane potential

To explore roles of *I_K1_* and hyperpolarization-activated Na^+^/K^+^ current (also known as the funny current), *I_h_*, in the elevated resting membrane potential of hiPSC-CM (again iCell1), we modulated *I_K1_* currents in the hiPSC-CM model. While keeping *I_h_* constant, increasing *I_K1_* current from zero or a low level indicated by expression data in hiPSC-CMs to that of an hAVCM lowered the elevated resting membrane potential back to ~-88mV (Figure 5**a**). Changing the conductance of *I_h_* on the other hand, had little effect on changing the AP regardless of the *I_K1_* expression levels (*g_K1_/g_K1,0_*= 1 or 0, Figure 5**b, c**). After restoring the resting membrane potential by elevating *I_K1_*, the simulation shows that *I_Na_* becomes a more active player in depolarizing the hiPSC-CM’s membrane potential.

**Figure 5.**
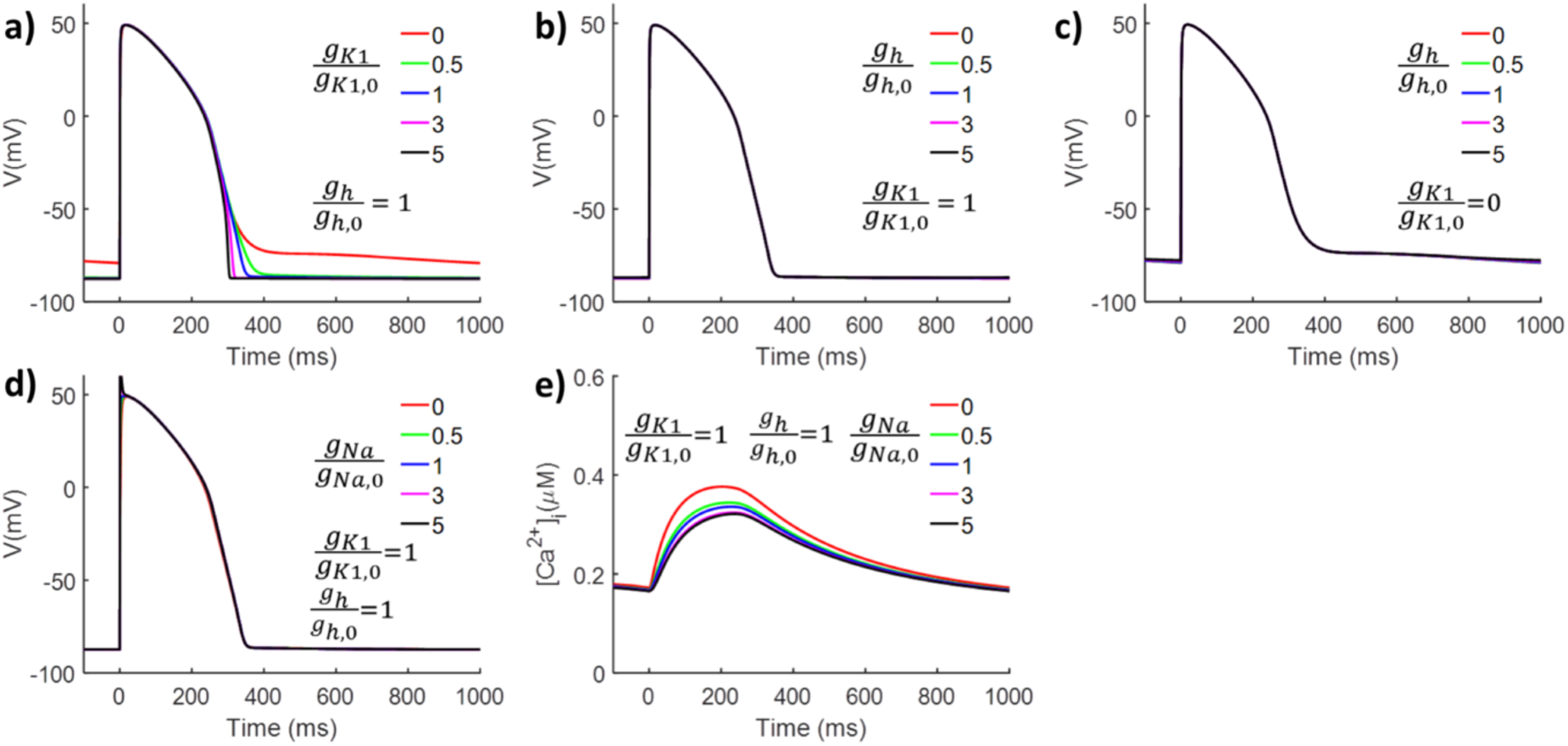
Influence of inward rectifier K^+^ and hyperpolarization-activated Na^+^/K^+^ channel conductance on the iCell1 version of the hiPSC-CM model. While keeping conductances of hyperpolarization-activated Na^+^/K^+^ channel, *g_h_,* at the level of hAVCM, increasing conductance of inward rectifier K^+^channel, *g_K1_,* lowered the resting membrane potential to ~ −90mV (**a**). If *g_K1_* was kept at the level of hAVCM, changing, *g_h_*, had no effects on action potential profiles (**b**). If *g_K1_* is low, changing *g_h_* had no effects to lower the resting membrane potential (**c**). When *g_K1_* and *g_h_* are at the level of hAVCM, effects of changing fast Na^+^ channel conductance, *g*_*Na*_, on action potential profiles (**d**) and calcium transients (**e**) became similar to those of hAVCM (Figure 4 **e** and **g**).

### Reduced contribution of sarcoplasmic reticulum (SR) in Ca^2+^ transients of hiPSC-CMs

To see the functional difference in calcium induced calcium release from the SR in hiPSC-CMs, we varied the flux through the ryanodine receptor (RyR) and sarcoplasmic/endoplasmic reticulum Ca^2+^ ATPase (SERCA) in both models. Both independent and concurrent changes in flux of RyR and SERCA had little effect on AP profiles (Figure 6**a, b, c**). CaT showed little change when only RyR flux was varied (Figure 6**d**) and moderate change when only SERCA flux was varied (Figure 6**e**). Combined effects of changing RyR and SERCA had large effects on CaT (Figure 6**f**). In the hiPSC-CM simulation of Figure 4**d** at a *g_CaL_/g_caL,0_* = 5 we see a maximum amplitude of ~0.35 μM which is similar to the maximal amplitudes in Figure 6**d** however bringing SERCA flux up to hAVCM levels independently (Figure 6**e**) or concurrently with RyR flux (Figure 6**f**) results in cytosolic Ca^2+^ concentrations of about 0.5 and 1.25 μM respectively which is much higher than levels predicted in the hAVCM by the ORd model (Figure 4**c** at *g_CaL_/g_CaL,0_* = 1).

**Figure 6.**
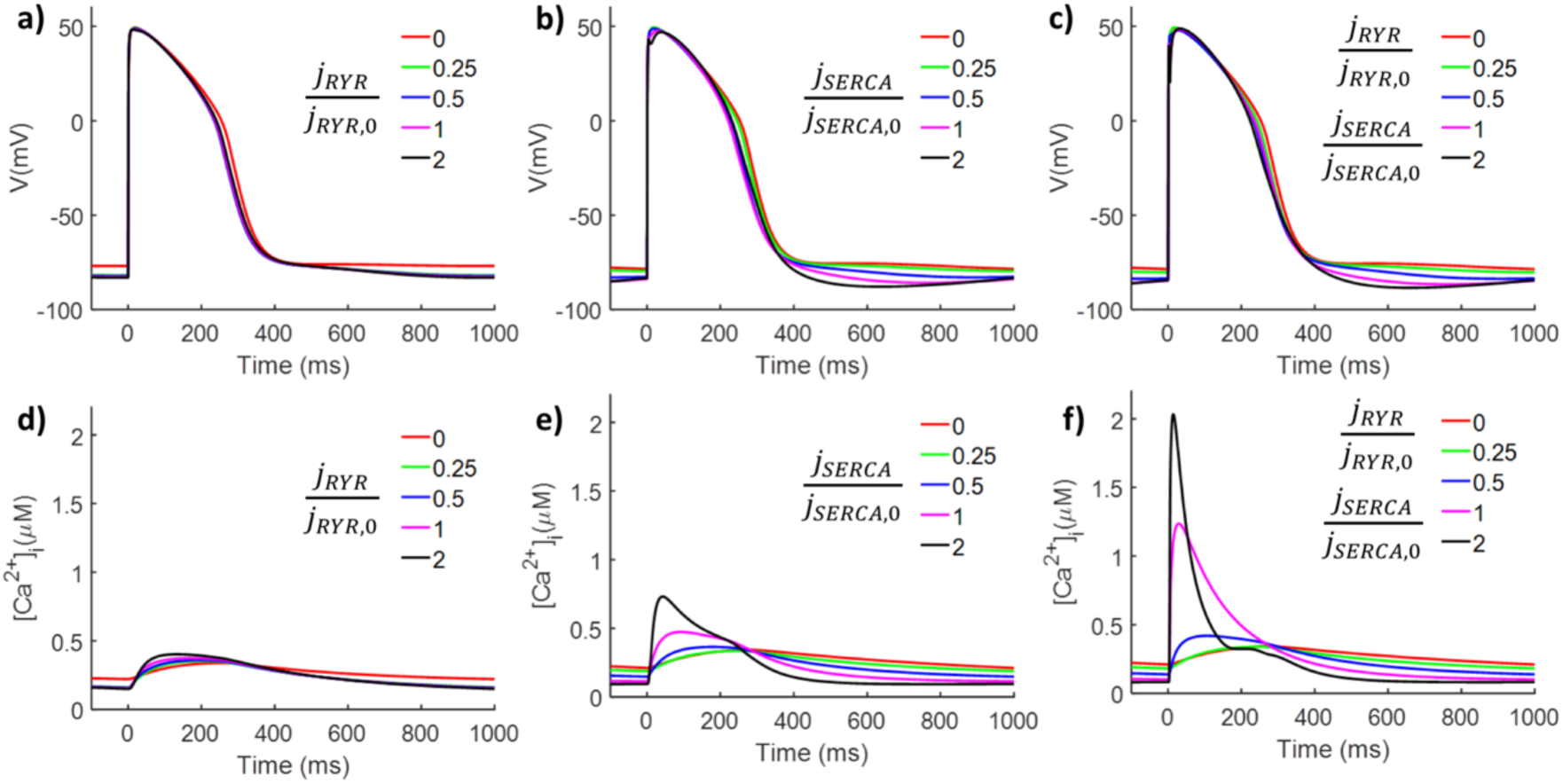
Effects of changing hiPSC-CM’s (iCell1) Ca^2+^release via ryanodine receptors, *j_RYR_,* and Ca^2+^ uptake via SERCA pump, *j_SERCA_*, on the simulated action potential profiles (**a**, **b**) and calcium transients (**d**, **e**). Increasing *j_RYR_* and *j_SERCA_* at the same time to the level of hAVCM reduced resting membrane potentials (**c**) and increased calcium transient’s amplitudes (**f**).

### hAVCM and hiPSC-CM proarrhythmic risk assessment of cardiac drugs

To visualize the differences in proarrhythmic sensitivities between hAVCM and hiPSC-CM we theoretically blocked the rapid delayed rectifier K^+^, fast Na^+^ (including the slow Na^+^ component) and L-type Ca^2+^ channels from 0 to 100% in different combinations (Figure 7). Blocking of each channel was accomplished by scaling the conductance of that channel between 0 to 1 (1 being normal conductance and 0 being a complete block). Each set of blocking parameters defined a position in the K^+^, Na^+^ and Ca^2+^ space and the APD_90_ of the resulting AP simulation was calculate at each position. APD_90_ values were scaled across a blue to red spectrum and we added separate colors for: arrhythmia including EAD (purple) and unstable AP or no depolarization (black). The 3D plots revealed that hiPSC-CM are more susceptible to *I_Kr_* and *I_CaL_* block and exhibit larger regions in the 3D space of arrhythmia as indicated in purple and black at both 1 Hz (Figure 7**a, b**) and at 0.5 Hz (Figure 7c, d). The interplay between *I_Kr_* and *I_CaL_* block describes much of the effects seen as is observed when we project the entire 3D space onto a K^+^ and Ca^2+^ plane (Figure 7**e-h**). 2D cross-sectional plots of *I_Na_* and *I_CaL_* block keeping *I_Kr_* constant shows no effect on APD_90_ as a function of *I_Na_* block (Supplement Figure S1).

**Figure 7.**
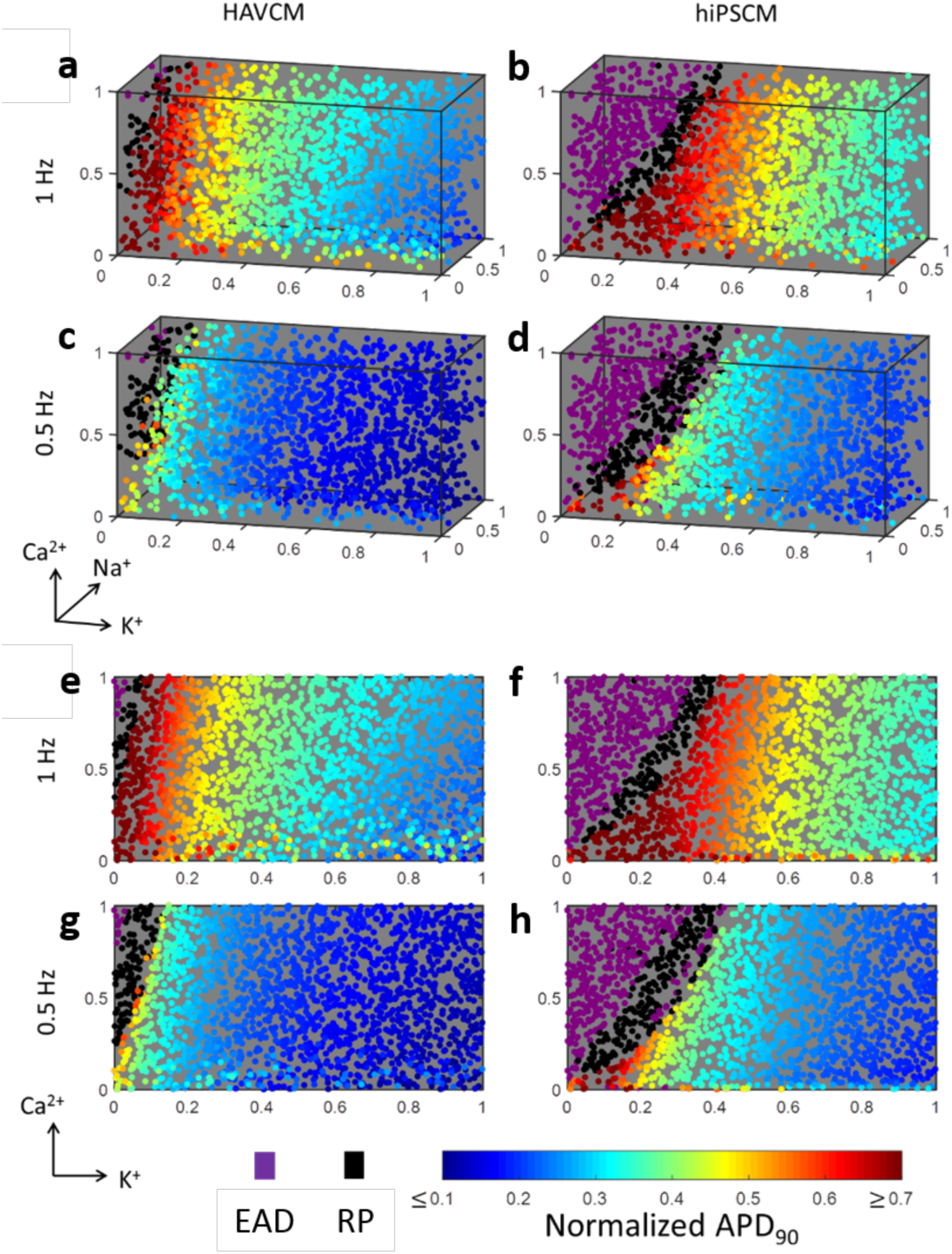
Action potential durations at 90% from their peak (APD_90_) were normalized by pacing period (i.e., 1s and 2s for 1 and 0.5 Hz pacing frequency, respectively) and color-coded (blue – red). Proarrhythmic action potential profiles were categorized into those exhibiting early after depolarization (EAD, purple) and repeating oscillation (RP, black). Combined effects of inhibiting conductance of K^+^, Na^+^, and Ca^2+^ channels at various degrees were plotted in 3D space with the color code for hAVCM (**a**) and hiPSC-CM (iCell1) (**b**) that were electrically paced at 1Hz. Decreasing pacing frequency to 0.5 Hz enlarged a region exhibiting EAD and RP in hAVCM (**c**) and hiPSC-CM (**d**). 2D scatter plots of hAVCM (e and g) and hiPSC-CM (f and h) were generated by projecting color-coded dots on to a K^+^-Ca^2+^ plane. Latin hypercube sampling was used to select combinations of K^+^, Na^+^, and Ca^2+^ inhibition.

Analyzing projections of 3D plots onto a 2D plane defined by *I_Kr_* and *I_CaL_* axis (Figure 7**e-h**) further clarified that an *I_CaL_* inhibition can mitigate APD_90_ elongation and/or arrhythmic AP profiles induced by *I_Kr_* blocking especially in hiPSC-CMs. The simulations also show that reducing the pacing frequencies enlarged arrhythmia sensitivity zone (purple and black) in both hAVCM and hiPSC-CM. At the lower pacing frequency (Figure 6**g, h**), an increased *I_Kr_* block moves AP profiles from moderate APD_90_ elongations rapidly into regions of arrhythmia sensitivity zone skipping over extremely elongated APD_90_ zones indicated by red and orange.

## Discussion

In this study, previous measurements of relative gene expression of key ion channels, exchangers, pumps and receptors between hAVCM and hiPSC-CM cell lines^7^ has been used to modify an existing mathematical model of the hAVCM (ORd model^14^) resulting in a novel hiPSC-CM electrophysiology and ion handling model. By using relative expression data which can be obtained across different cell lines and even batches of a given cell line we are able to develop a theoretical hiPSC-CM model that qualitatively represents the variation in cell function and can be matched to a specific experiment preparation. This expression-based hiPSC-CM model not only exhibited previously observed characteristic differences between hiPSC-CMs and hAVCMs, but qualitatively reproduced the trends in APD_90_ observed across the 3 different batches of 2 different cell lines by Huo et al.^7^ (Table 2). The experimentally observed characteristic differences reported from various laboratories have been summarized previously by Hoekstra et al.^16^ and include 1) an elevated resting membrane potential in an hiPSC-CM (−57 to −75mV) when compared to an hAVCM (−81.8 to −87mV), 2) a reduced maximal upstroke velocity, (dV/dt)_max_, in hiPSC-CMs, 3) a longer APD_90_ in hiPSC-CM than hAVCM and 4) spontaneously beating of hiPSC-CMs. Our expression based hiPSC-CM model exhibits the first three observed characteristics except in the case of modeling the Cor4U lines where both model and observation show a shorter APD_90_ in contradiction with the third observation by Hoekstra et al.^16^ The expression based hiPSC-CM model does not exhibit spontaneous beating, also known as automaticity, however this is likely because we have removed the ohmic-like Na^+^ background leak current in the model which are typically used to drive resting membrane depolarization and spontaneous beating frequency.

**Table 2.** Beating rate (BR) and field potential duration (FPD) for iCell and Cor4U hiPSC-CM lines from Huo et al.^7^.

|  | iCell 1 | iCell 2 | iCell 3 | Cor4U 1 | Cor4U 2 | Cor4U 3 |
| --- | --- | --- | --- | --- | --- | --- |
| BR (bpm) | $36.1 \pm 1.3$ | $35.8 \pm 1.3$ | $36.7 \pm 1.5$ | $50.9 \pm 7.2$ | $67.3 \pm 4.2$ | $82.2 \pm 3.9$ |
| FPD (ms) | $524.0 \pm 24.7$ | $522.5 \pm 40.5$ | $455.6 \pm 29.0$ | $415.3 \pm 57.4$ | $256.6 \pm 29.8$ | $172.2 \pm 23.9$ |

This new expression based hiPSC-CM model has distinct advantages over the most recent hiPSC-CM model by Paci et al.^12^ which is based on patch clamp data from Ma et al.^13^. First and foremost, this model can be developed from expression data alone and does not rely on expensive and time-consuming patch clamp data for each critical ion channel driving hiPSC-CM electrophysiology. Expression data can be tractably obtained from multiple cell lines and therefore can reflect variation in hiPSC-CM function across different cell lines or at different maturation time points. This opens the door to parameterizing hiPSC-CM models which capture the variability needed to reflect patient specific function. Another point of note is that we have eliminated the background leak currents of Na^+^ and K^+^ currents in the model and replaced them with the ultra-rapid K^+^ and hyperpolarization activated Na^+^/K^+^ channel currents (*I_Kur_* and *I_h_).* The *I_Kur_* and *I_h_* currents can be quantified through relative expression and therefore this hiPSC-CM model does not rely on unidentifiable leak currents to drive resting membrane depolarization and spontaneous beating. The Paci et al. model^12^ relied on a large Ca^2+^ leak current to drive resting membrane potential depolarization to represent automaticity which is not used here.

This hiPSC-CM model showed a slowly depolarizing diastolic membrane potential resulting in an elevated resting membrane potential that was especially evident at slow pacing frequencies. A higher resting membrane potential reduces the functional availability of fast Na^+^ channels since the inactivation gate cannot fully reset and the reduction in Na^+^ influx is further reduced by the lower current density of the fast Na^+^ channel as indicated by reduced SCN5A expression across all batches of the two hiPSC-CM lines. Shifting the major contributor of CM depolarization from fast *I_Na_* to relatively slower *I_CaL_* explains the lowered dV/dt^max^ in hiPSC-CMs. Analysis of the simulations shown in Figure 5 identified that a significantly reduced expression of *I_K1_* is the primary cause of slowly elevating resting membrane potential as demonstrated experimentally by reintroducing the *I_K1_* channel through forced expression in human embryonic stem cell-derived cardiomyocytes^11^ and electrophysiologically in hiPSC-CMs^11,17^.

This expression based hiPSC-CM model is also able to quantitatively capture the relatively immature sarcoplasmic reticulum (SR) phenotype known to exist in hiPSC-CMs^18^. Previously reported electron micrographic observations of the immature SR showing a lack of T-tubules in hiPSC-CMs^19^ that belies the significantly reduced expression of RyR and SERCA. This immature phenotype is believed to play a role in the longer APD_90_ typically observed in hiPSC-CMs but longer APD_90_ also involves the interplay between the increased L-type Ca^2+^ channel and relatively unchanged rapid delayed rectifier K^+^ channel currents as reflected in the expression data (Table 1). Increased *I_CaL_* conductance results in a higher maximum L-type Ca^2+^ currents in hiPSC-CMs; however, the reduced release and uptake of Ca^2+^ through the RyR and SERCA pump, respectively, leaves in question the ultimate profile of the calcium transient (CaT) for each cycle. The simulations for a normal hAVCM shown in Figure 4**c** at *g_CaL_/g_CaL,0_* = 1 and the iCell1 shown in Figure 4**d** at *g_CaL_/g_CaL,0_* = 5 indicate that the peak cytosolic Ca^2+^ for the hiPSC-CM is actually higher and decays more slowly than in the hAVCM. However, reduced RyR in hiPSC-CMs results in a sensitivity of cytosolic Ca^2+^ to changes in *g_CaL_* that was smaller than in hAVCMs. While the SR in hAVCMs efficiently removed an elevated intracellular Ca^2+^ in diastole, the capacity of hiPSC-CMs to remove intracellular Ca^2+^ during diastole was compromised. This is due to the reduced ability of the SR in hiPSC-CMs to take up intracellular Ca^2+^ through SERCA.

Calcium ion is known to be the mediating signaling molecule responsible for translating action potential into cardiac contraction. The gene expression data (Table 1 and Huo et al.^7^) reveals that in hiPSC-CMs both RyR and SERCA pump are expressed at a much lower level when compared to hAVCMs. Therefore, the calcium ions in the cytosol enter mainly through *I_CaL_* which cannot produce the rapid change in Ca^2+^ usually observed in hAVCMs. This immature SR phenotype is similar to that observed in CMs during cardiac development^18^.

Simulations of hiPSC-CM (iCell1) over a range of fast Na^+^ channel conductances (Figure 4**h**) and inward rectifier K^+^ conductance (implied by Figure 5**e**) also show little sensitivity on cytosolic Ca^2+^. Independently increasing the RyR in the hiPSC-CM to baseline hAVCM levels makes little difference in the CaT and independently increasing the SERCA flux shows a small change in CaT. However, increasing both concurrently in the hiPSC-CM makes a dramatic difference in CaT. Since the CaT will be important to predict accurately as we measure and analyze force generation in hiPSC-CMs, understanding how each ion channel, pump and receptor involved in the composition of the CaT will be important.

While we do see a depolarized resting membrane potential in our expression-based hiPSC-CM model, the membrane potential does not rise to the threshold (-40mV) needed to spontaneously fire. There are two explanations for this lack of automaticity in the model. First, the combination of the absence of the Na^+^ and K^+^ background leak currents, which have been eliminated in this model, and the inclusion of *I_h_*, which is the major depolarizing current used to replace the omitted background currents are not able to generate a constant depolarization as the membrane potential drifts away from the typical resting membrane potential of ~ −88 mV. Automaticity could be achieved by inserting a small background Na^+^ or changing the conductance of the retained background Ca^2+^, however full AP time courses would be necessary to identify the background channel conductance needed to achieve automaticity. The second reason is that automaticity may be driven by a single or small group of cells that have enough depolarization to generate an AP. AP generation from this region then would drive depolarization in neighboring hiPSC-CMs propagating across the entire tissue. Observations of CaT propagation in a two-dimensional space (8x8 mm^2^) made in our lab (result not shown) show that CaT was initiated from a single site and rarely from more than that one site. The pacemaker-like hiPSC-CMs may also express T-type Ca channel that can be depolarized at lower membrane potential and drive automaticity but would not show up as significantly present in our expression data.

The strength of this theoretical approach is that we are now able to visualize the proarrhythmia potential of drugs that block the typical cardiomyocyte ion channel targets such as fast Na^+^, L-type Ca^2+^ and rapid delayed rectifier K^+^ channels in both a normal hAVCM and a representative hiPSC-CM. Figure 7, where the hiPSC-CM is represented by iCell1, shows: 1) hiPSC-CMs are predicted to have a higher proarrhythmia risk in response to ion channel blockage than hAVCMs 2) proarrhythmia risk associated with *I_Kr_* blockage can be mitigated by concurrent blocking of *I_CaL_* and 3) blocking of fast Na^+^ channels has little effect on proarrhythmia risk (also see Supplement Figure S1). In addition, we are able with this methodology to understand why different hiPSC-CM cell lines might actually show different responses. iCell lines would be posited here to exhibit a higher proarrhythmia risk than Cor4U cell lines since the iCell lines exhibit a longer APD_90_ than Cor4U even before a theoretical blocking agent is applied.

## Limitations

In this study, we are using relative expression data of each hiPSC-CM cell line and batch with respect to expression levels to hAVCM. We are assuming that the hAVCM expression levels represent the conductances, permeabilities and maximal fluxes given in the ORd model. This assumption does not affect the qualitative observations we are discussing in this study. However, to match AP and CaT time courses along with generating a model that spontaneously beats at the correct frequency, the reference hAVCM model must be identified with a series of experiments similar to that proposed by Krogh-Madsen et al.^20^.

The Ca^2+^ release kinetics through RyR in the original ORd model depends on the conductance of *I_CaL_,* SR junctional space Ca^2+^ concentration, and activity of CaMKII^21^. This description is representative of SR function in hAVCMs where the dyadic space formed between T-tubule and sarcoplasmic reticulum (SR) and the pairing of RyR and L-type calcium channel is generally accepted. This structural description, however, may not be present in hiPSC-CMs that have significantly reduced RyR and SERCA expression and are thought to be not as tightly coupled to the release of Ca^2+^ through L-type Ca^2+^ channels in the dyadic space. As more detail becomes known about the structural and functional details of the dyadic space and SR function in hiPSC-CMs, we would be able to modify SR function based on more than just RyR and SERCA expression.

## Conclusion

We have modified a widely accepted model for the electrophysiology and ion handling in an hAVCM^14^ using relative expression data of ion channel, exchange, pump and receptor protein encoded genes to develop a novel model of the hiPSC-CM. This model can represent variability in hiPSC-CM function across different cell lines and cell line batches to qualitatively represent difference in action potential duration. The hAVCM model used in this development is currently supported by CiPA for the testing of proarrhythmia risk^22^. This expression-based hiPSC-CM model suggests that using hiPSC-CMs for testing of proarrhythmia risk of drug compounds over predicts the risk for K^+^ and Ca^2+^ ion channel blocking compounds.

## Methods

### Theoretical model of human induced stem cell derived cardiomyocytes

This study modified the ORd hAVCM model^14^ to reflect the relative differences in the expression of eight different transmembrane ion channels, the Na^+^/Ca^+^ exchanger along with ryanodine receptor and the calcium pump in the SR in hiPSC-CMs against those in hAVCMs^7^. The ORd model is a comprehensive electrophysiology and ion handling model that was developed based on experimental measurements on dynamic properties of ion channels and pumps using isolated healthy hAVCMs. The relative expression data is then used to scale the conductance of, permeability of, or flux through each specific channel, exchanger, pump or receptor in the theoretical model creating an expression-based hiPSC-CM model (see example set of equations in the Supplement). It is assumed that there is no change in the gating or binding kinetics in any of the selected transport mechanisms and the number of fully functional channels and transporters in cells is a liner function of gene expression.

The gene expression analysis of hiPSC-CMs^7^ detected ultra-rapid K^+^ (Kv1.5, KCNA5) channels and hyperpolarization-activated cation (HCN2 and HCN4) that are not represented in the ORd model. The Kv1.5 channel protein has been detected at equal levels in human atrial and ventricular cardiomyocytes^23^ however the ultra-rapid K^+^ current (*I_Kur_*) is either non-existent or reduced in hAVCMs^24^. Similarly, the gene expression of HCN2 and HCN4 is seen in both atrial (specifically the sinoatrial node (SAN) and ventricular tissue, however the ratio of HCN4/HCN2 is much larger in the SAN than in the ventricle. In addition, the total amount of HCN has been reported to be as much as 7.5 times lower in ventricular cardiomyocytes than in SAN cells, suggesting a reduced current density in hAVCMs.

To represent these two missing channel currents in the ORd model we looked at previous modeling studies of atrial and ventricular myocyte electrophysiology. The ultra-rapid delayed rectifier K^+^ current was formulated based on the human atrial cell electrophysiology model by Cortmanche et al.^25^ with the maximal conductance reduced about 6-fold from values found in human atrial cells as indicated experimentally^24^. For the hyperpolarization-activated cation current, *I_h_*, we used the model from Kurata et al.^26^ for the HCN2 channel based on data from Moroni et al.^27^. The advantage of using the Kurata et al. model is that instead of lumping the inflow current from Na^+^ and outflow current from K^+^ into a net current, each ionic current is represented separately. While both HCN2 and HCN4 are present in ventricular cardiomyocytes it has been shown in mouse ventricular tissue that over 75% of the HCN expression is that of the HCN2 isoform^28^. However, expression data from Huo et al.^7^ shows the opposite trend in hAVCM cells with 75% of the HCN expression in the form of HCN4.

The major functional difference between the four different isoforms in the HCN family is their rate of activation with HCN1 having the most rapid and HCN4 having the slowest gating kinetics^29^. Even though there is some faster gating HCN2 in ventricular cardiomyocytes, measurements of the gating time constant, *τ_h_*, in human embryonic stem cell derived cardiomyocytes (heSCMs) is on the order of 2000 ms in the range of −110 to −90 mV^30^. This time constant is closer to the gating kinetics of the HCN4 isoform so we replaced the Kurata et al. gating kinetics with that used in the Paci et al. hiPSC-CM model ^12^. Because these two ion-channel currents (*I_Kur_* and *I_h_*) were not represented in the ORd model instead of adding them as additional currents we used them to replace the background Na^+^ and K^+^ channels in the original ORd model. The Ca^2+^ background channel current was retained. The maximal conductances of these two channels in the hAVCM led to currents that were close to the ORd background Na^+^ and K^+^ currents in hAVCMs and when changed as a ratio of expression to represent the hiPSC-CM cell lines resulted in an AP phenotype in diastole like that observed experimentally. Replacement of the two background channels with ion channels with conductance fixed to the expression data limited our ability to generate automaticity by changing conductance; however, a depolarized resting membrane potential was achieved with the expression-based modification of model conductances alone.

The dyad space was retained in the ORd model even though there is evidence that the formation of t-tubules is reduced in hiPSC-CMs. This will not necessarily affect the shape of the action potentials in the models used here but must be considered when sarcomere force generation is simulated from the cytosolic calcium transients. A representation of the modified ORd/hiPSC-CM model and the modifications made are shown in Figure 1.

An important feature of the ORd model is that given appropriate initial conditions the model goes to a steady state with stable APs as well as cytosolic concentrations of Na^+^, Ca^2+^ and K^+^ ([Na^+^]_i_, [Ca^2+^]_i_ and [K^+^]_i_, respectively). To check that this remained the case with the new inserted *I_Kur_* and *I_h_* for the hAVCM and all six hiPSC-CM parameterized model simulations we calculated the change at 10 different time points of V_m_, [Na^+^]_i_, [Ca^2+^]_i_ and [K^+^]_i_ between two successive APs. A steady state was reached in all cases with the error in each of the four variables dropping below a 0.1% change.

Simulations were performed for three batches from two different hiPSC-CM lines along with the hAVCM at stimulated at frequencies of 1 and 0.5 Hz. The stimulation was in the form of a step function of an applied voltage with magnitude −80 mV for 1 ms. From these six different representations of an hiPSC-CM, we chose to look at a single representative hiPSC-CM (iCell1) and compare it to the simulated response of the hAVCM. All simulations reached a steady state according to the criteria previously mentioned. All simulations were coded and run in Matlab (Matlab, Inc., Natick, MA, USA).

### Expression data and experimental methods

Expression data was obtained from researchers at the FDA and methods to obtain expression for the 3 batches of 2 different cell lines are described in detail in the study by Huo et al. ^7^. Briefly, three batches of hiPSC-CMs from two vendors and one hAVCM line were prepared according to the vendors specifications. The iCell cells were obtained from Cellular Dynamics International (Madison, Wisconsin), the Cor4U cells were obtained from Axiogenesis (Nattermannalee, Germany) and the hAVCMs were obtained from Biochain (San Francisco, California). Total RNAs were isolated at day 15 for the iCells and day 9 for the Cor4U cells and real-time qRT-PCR was performed on the cells to obtain expression of 16 cardiac genes related to the ion channels, exchangers, pumps and receptors used to quantify conductances and fluxes used here. The qRT-PCR expression values for the hAVCM were used to normalize the hiPSC-CM expression data. In addition, observations from the study of Huo et al^7^ of field potential duration (FPD) were compared to our simulations results for APD_90_ to confirm trends.

### Disclaimer

The views presented in this article do not necessarily reflect those of the Food and Drug Administration. The authors declare that there are no conflicts of interest regarding the publication of this article.

## Determining dV/dt^max^ from simulation results

When evaluating the slope of the voltage upstroke at the beginning of the action potential (dV/dt^max^) from simulation data, the value is highly dependent on the resolution of the time step selected. This means that when looking at experimental measurements of dV/dt^max^ the value can also vary depending on sampling rate. The table below shows at 1Hz a 19% to 88% increase in the calculated dV/dt^max^ when dividing the time from stimulation to peak voltage over 500 intervals to find the largest value as opposed to 50 intervals. It is often not clear what the sampling rate is for dV/dt^max^ obtained experimentally so we assumed the upstroke was divided into 100 intervals in this study yielding values of 342 mV/ms for the hAVCM and 134 mV/ms for iCell1.

**Table S1:** Comparison of model predicted (dV/dt)_max_.

| Frequency (Hz) | Sampling Number (N)* | $(dV/dt)_{\max}$ (mV/ms) | | | | | | |
| --- | --- | --- | --- | --- | --- | --- | --- | --- |
|  |  | Cor4U1 | Cor4U2 | Cor4U3 | hAVCM | iCell1 | iCell2 | iCell3 |
| 1.0 | 50 | 96 | 88 | 103 | 297 | 102 | 100 | 109 |
|  | 100 | 112 | 121 | 114 | 342 | 134 | 120 | 153 |
|  | 300 | 149 | 144 | 129 | 354 | 167 | 166 | 198 |
|  | 500 | 154 | 146 | 130 | 354 | 172 | 174 | 206 |
| 0.5 | 50 | 83 | 82 | 82 | 311 | 84 | 84 | 84 |
|  | 100 | 86 | 84 | 84 | 342 | 88 | 87 | 89 |
|  | 300 | 89 | 87 | 85 | 355 | 93 | 92 | 97 |
|  | 500 | 89 | 87 | 85 | 356 | 94 | 93 | 98 |

\* Dividing the upstroke phase of action potential into N equal intervals to calculate $dV/dt$ .

Supplemental Table S1. Variation in calculated dV/dt^max^ from simulation as a function of how the time from AP initiation to peak is divided. More time divisions (higher sampling number) capture the steepest section of the AP upstroke without averaging it over less steep regions. dV/dt^max^ measures obtained experimentally appear to be sampled at rates corresponding to a sampling number of 50-100 in this table.

## Variation of APD_90_ with respect to fast Na^+^ block for iCell1

To show how invariant APD_90_ is to Na^+^ block we took slices spanning a 0.1 range of K^+^ block of our Na^+^/K^+^/Ca^2+^ blocking simulations for both hAVCM and hiPSC-CM (iCell1) at 1 Hz and 0.5 Hz. All simulations contained within that slice are then projected onto the Na^+^/Ca^2+^ block plane to render the images below. It can be seen that at horizontal lines (constant Ca^2+^ and varying Na^+^ block) across each one of these slices shows little variation in color indicating little change in APD_90_. This is to be expected in the hiPSC-CMs since in each hiPSC-CM model simulated the resting membrane potential is sightly depoarized (−80 to −70 mV) preventing the inactivation gate of the fast Na^+^ channel from resetting so block will have little effect on AP shape and APD_90_. In addition, the expression of fast Na^+^ (SCN5A) is reduced to 0.65 of the expression of the hAVCM further limiting the effects of a fast Na^+^ block. Even though the variation is larger in hAVCMs for fast Na^+^ block it still shows a minimal contribution compared to L-type Ca^2+^ block.

**Supplement Figure S1.**
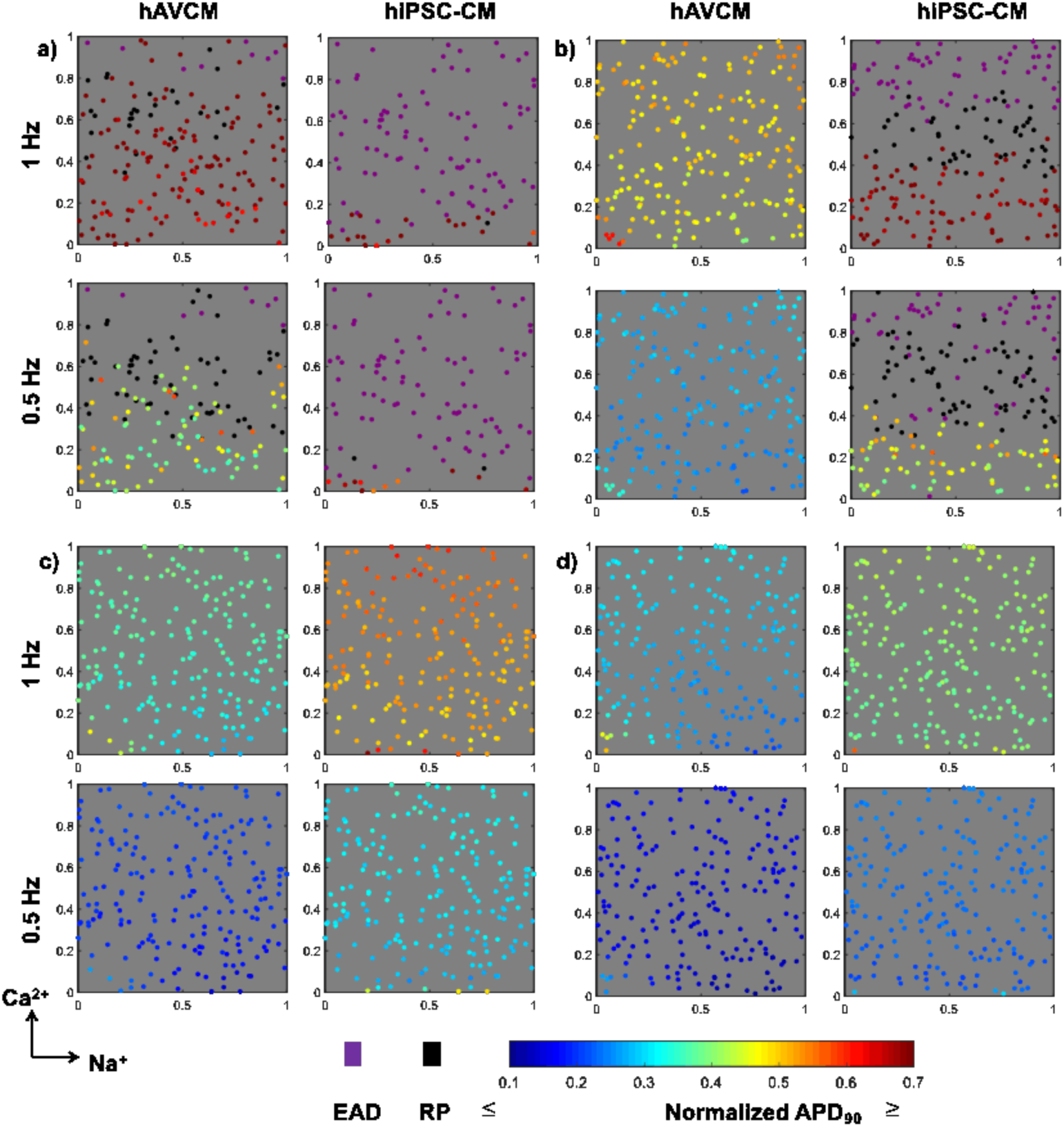
2D scatter plots by projecting color-coded dots of 3D scatter plots in Figure 7**a-d** to a Na^+^/Ca^2+^ planes with the K^+^ axis ranging from a) 0 – 0.1, b) 0.2 – 0.3, c) 0.45 – 0.55 and d) 0.7 – 0.8.

## Example of expression based scaling of ion channel, exchanger, pump and receptor function

Each of the ion channels, exchangers, pumps and receptors for which expression data was obtained are modified to reflect the relative expression ratio with respect to the expression found in the hAVCM line. The modification assumes that the kinetic parameters and single component function remains the same and expression solely reflects the relative number of fully functioning ion channels, transporters, etc. This means that in the formulation of an ion channel we are purely scaling the maximal conductance parameter by the expression ratio. The maximal conductance represents the conductance if all channels of that particular type on the cell membrane were open. Maximal conductance parameters were scaled for the: fast Na^+^ channel, transient outward K^+^ channel, rapid delayed rectifier K^+^ channel, slow delayed rectifier K^+^ channel, inward rectifier K^+^ channel, Na^+^/Ca^2+^ exchanger, hyperpolarization activated Na^+^/K^+^ channel and ultrarapid K^+^ channel. For the L-type Ca^2+^ channel the maximal permeability was scaled and for the ryanodine receptor and sarco/endoplasmic reticulum Ca^2+^ ATPase maximal fluxes were scaled.

As an example of how we altered the original ORd formulation, the complete formulation for the delayed rectifier K^+^ channel is given below with the modification due to expression data in bold font:

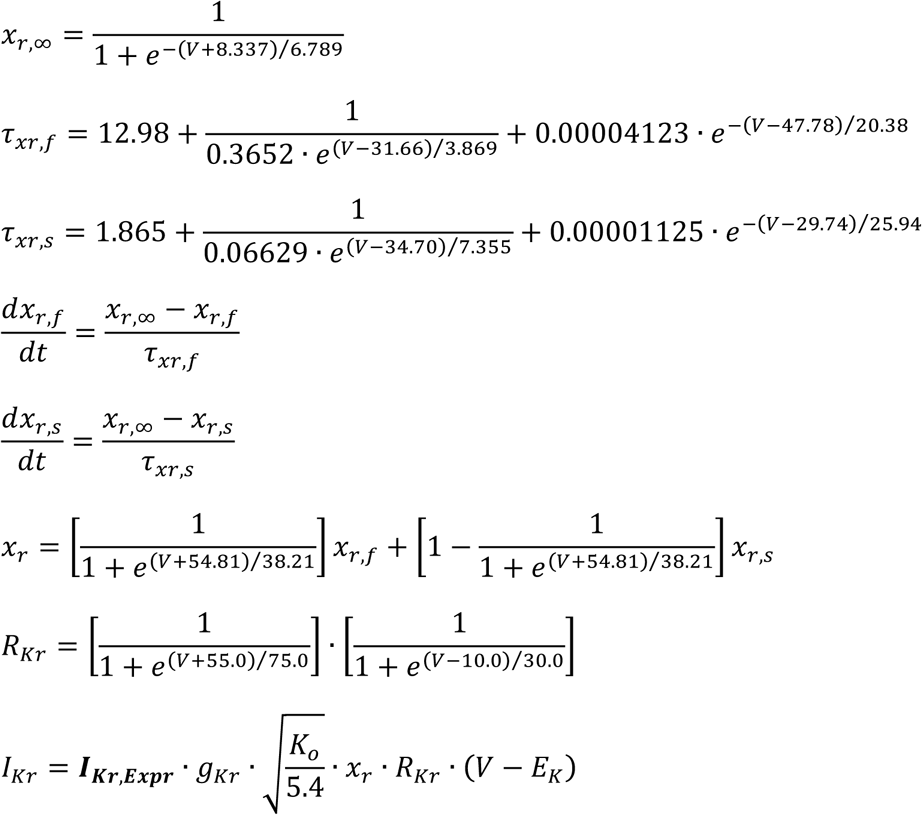

Where *x*_*r*,∞_ is the steady state open probability of the activation gate, *V* is the membrane potential, τ_*xr,f*_ isthe time constant of the fast activation component of the activation gate, τ_*xr,s*_ is the time constant of the slow deactivation component of the activation gate, *x_r_* is the open probability of the activation gate, *R_Kr_* is the instantaneous open probability of the inactivation gate, *l_Kr_* is the current through the rapid delayed rectifier channel, *I_Kr,Expr_* is the ratio of the KCNH2 expression in the hiPSC-CM with respect to that measured in the hAVCM, *g_Kr_* is the maximal rapid delayed rectifier conductance of an hAVCM, *K_o_* is the extracellular K^+^ concentration and *E_K_* is the Nernst potential for K^+^.

In three of the channels (fast Na^+^, L-type Ca^2+^ and rapid delayed rectifier K^+^) blocking is achieved by adding an additional term to the equation for current. Therefore, in the case of the rapid delayed rectifier current given above *I_Kr_* is given by:

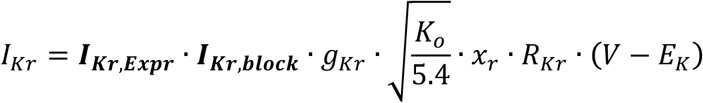

where *I_Kr,block_* is the fraction of total current flowing through the channel after block and goes from 0 (100% block) to 1 (no block).

